# Concurrent infection with *Mycobacterium tuberculosis* confers robust protection against secondary infection in macaques

**DOI:** 10.1101/403691

**Authors:** Anthony M. Cadena, Forrest F. Hopkins, Pauline Maiello, Allison F. Carey, Eileen A. Wong, Constance J. Martin, Hannah P. Gideon, Robert M. DiFazio, Peter Andersen, Philana Ling Lin, Sarah M. Fortune, JoAnne L. Flynn

**Affiliations:** Department of Microbiology and Molecular Genetics, University of Pittsburgh School of Medicine, Pittsburgh, Pennsylvania, USA; Department of Immunology and Infectious Diseases, Harvard T.H. Chan School of Public Health, Boston, Massachusetts, USA; Department of Pathology, Massachusetts General Hospital, Boston, Massachusetts; Statens Serum Institut, Copenhagen, Denmark; Department of Pediatrics, Children’s Hospital of Pittsburgh, University of Pittsburgh Medical Center, Pittsburgh, Pennsylvania, USA; Ragon Institute of Massachusetts General Hospital, Massachusetts Institute of Technology and Harvard, Cambridge, Massachusetts, USA

## Introduction

For many pathogens, including most targets of effective vaccines, infection elicits an immune response that confers significant protection against reinfection. There has been significant debate as to whether natural *M. tuberculosis* (Mtb) infection confers protection against reinfection. Here we experimentally assessed the protection conferred by concurrent Mtb infection in macaques, a robust experimental model of human tuberculosis (TB), using a combination of serial imaging and Mtb challenge strains differentiated by DNA identifiers. Strikingly, ongoing Mtb infection provided complete protection against establishment of secondary infection in over half of the macaques and allowed near sterilizing bacterial control for those in which a secondary infection was established. By contrast, boosted BCG vaccination reduced granuloma inflammation but had no impact on early granuloma bacterial burden. These findings are evidence of highly effective concomitant mycobacterial immunity in the lung, which may inform TB vaccine design and development.

## Main text

Epidemiologic studies suggest that primary *Mycobacterium tuberculosis* (Mtb) infection provides up to 80% protection against TB disease due to secondary exposure^1^, although assessing protection against actual reinfection is not possible. However, up to ∽20% of patients who complete drug treatment develop TB again, in part due to reinfection^2-6^. In mice, ongoing or treated Mtb infections only reduce bacterial burdens in the lung by ∽10-fold, roughly equivalent to BCG vaccination^7,8^. We sought to quantitatively assess the effect of a concurrent Mtb infection on rechallenge in the cynomolgus macaque model^9-11^, which recapitulates nearly all aspects of human Mtb infection. To probe the dynamics of reinfection, we used serial [^18^F]-fluorodeoxyglucose (FDG)-PET-CT imaging to track timing of granuloma formation following secondary exposure^12-14^, which allowed all granulomas to be retrieved at necropsy. The outcomes of primary and secondary challenges were defined with Mtb libraries marked with unique DNA identifiers that were tracked by sequencing and/or a custom direct hybridization (NanoString) assay^12,15,16^.

Eight macaques received a primary challenge with Mtb Erdman library A (<15 CFU) (**Fig. 1A**). As expected based on our published work^10,11,17^, this infection resulted in a range of outcomes by 16 weeks as assessed by PET-CT^11^, from minimal to progressive disease (**Fig. 1B, Supp. Table 1**). Sixteen weeks after primary infection, animals were rechallenged with Mtb Erdman library B (<15 CFU). Six naïve control animals were challenged with library B in parallel (**Fig. 1A**). Granuloma formation after library B challenge was tracked for ∽4 weeks by PET-CT imaging (**Supp. Table 1**) (**Fig. 1C**). The number of granulomas detectable by PET-CT imaging at 4 weeks post-infection in naïve animals is one correlate of the number of bacteria that successfully establish infection^12,13^. Individual granulomas and lymph nodes were obtained at necropsy (4 weeks post-library B) for bacterial and immunologic analyses. At this time point, which is just prior to the onset of adaptive immunity in naïve animals, granuloma bacterial burdens are relatively uniform and at their highest levels with minimal bacterial killing in naïve animals ^12,18,19^. Thus, comparing granulomas at 4 weeks post-library B in naïve and reinfected animals allows a direct comparison of early bacterial control.

**Figure 1:**
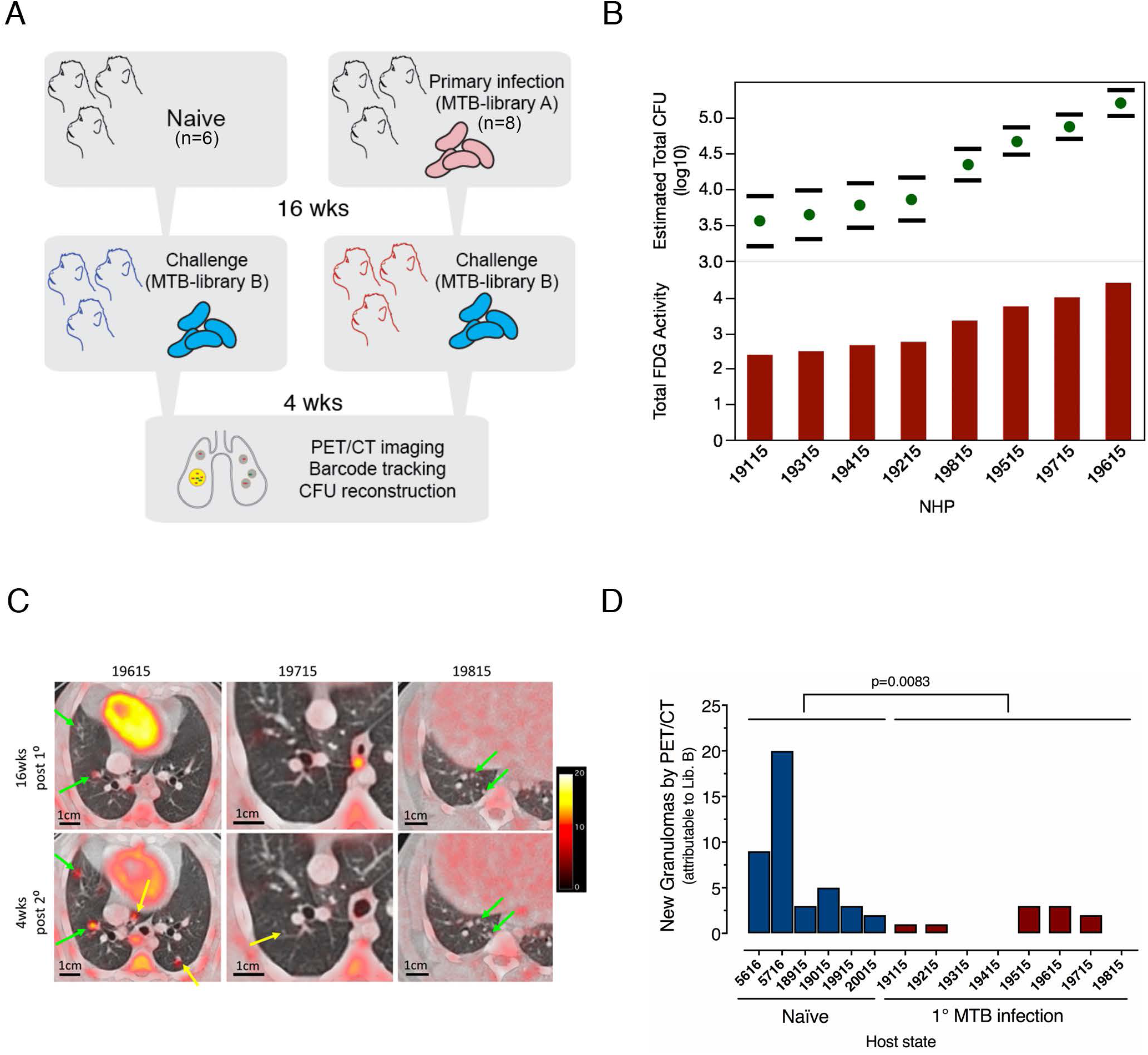
Concurrent Mtb infection limits the establishment of new granulomas. (a) Experimental schema of reinfection: Macaques (N=8) were infected for 16 weeks with low dose (<15 CFU) Mtb library A and then rechallenged with low dose Mtb library B for 4-5 more weeks. Naïve (N=6) macaques were challenged in parallel with only Mtb library B. (b) Low dose library A (primary) infection in the reinfection cohort resulted in a spectrum of host outcomes at 16 weeks as assessed by [^18^F]-FDG-PET/CT; total thoracic bacterial burden was estimated from the lung FDG activity and error bars denote 95% confidence intervals. (c) Imaging was used to discriminate primary strain infection and dissemination (green arrows), new granulomas after re-infection (yellow arrows); three different macaques are shown at 4 months post-primary infection, and 4 weeks post-reinfection. Monkey 19815 (far right panels) had no new granulomas detected after secondary infection. (d) Macaques with a primary infection had fewer newly established secondary granulomas seen by imaging and confirmed as containing Mtb Library B DNA at 4 weeks vs. naïve control monkeys (p=0.0083, Mann-Whitney test).

Using PET-CT to identify formation of new granulomas, there was no aggregate difference between new granuloma formation after library B challenge in the presence or absence of ongoing infection (**Supp. Figure 1A)**. However, this metric alone cannot distinguish new granuloma formation due to library B challenge from granulomas formed by ongoing dissemination from sites of primary infection (library A). Indeed, three animals had apparent dissemination just prior to or during secondary infection, based on imaging (monkey IDs: 19415, 19515, 19615). Moreover, imaging alone fails to capture the potential for library B seeding sites of existing infection as described in fish, frogs, and mice^20,21^. Therefore, we deconvoluted the identity of library tags in granulomas formed after primary and secondary infections. Tags were identifiable both from bacteria cultured from granulomas and, in most cases, from tissue homogenates including the homogenates of sterile lesions.

As expected, DNA tags from sampled granulomas in naïve animals mapped entirely to library B (**Fig 1D, Supp. Table 2**). In animals challenged during ongoing primary infection, there was no significant difference in total number of granulomas containing library B as compared to naïve animals (p=0.1525) (**Supp. Figure 1B**). Nine granulomas (of 95 total analyzed) from reinfected monkeys contained both library A and library B DNA (Supp. Table 2), suggesting that occasionally library B seeded pre-existing sites of infection. Therefore concurrent infection did reduce the number of *new* granulomas attributable only to library B (**Fig. 1D**, p=0.0083). The number of granulomas attributable only to library B at 4 weeks was also significantly lower than the number of granulomas established by library A granulomas at 4 weeks (as determined by PET CT) in the same animals (p=.0156, Supp. Fig. 1C). Indeed, there was complete protection in one animal with no detectable library B DNA in any tissues (monkey ID: 19815) (**Fig 1D**, **Supp. Fig. 1A, B)**.

To assess growth of the secondary challenge strain, we evaluated the bacterial burdens of individual granulomas in both cohorts and used the library tags to attribute these bacterial loads to library A and/or library B (**Fig. 2A**). In naïve animals, the distribution of granuloma bacterial burdens (library B) was consistent with our previous studies^12^ with a median bacterial load of 8300 and few sterile granulomas. By contrast, most granulomas from reinfected animals had lower bacterial loads than in naïve animals, with many sterile granulomas (**Fig. 2A,** p<0.0001). Strikingly, in concurrently infected animals, the few library B granulomas that did form had markedly lower bacterial burdens than in naïve macaques where these granuloma were assessed at the same time post-library B infection (**Fig. 2A, 3C**). In addition to the one animal that had no library B DNA (monkey ID: 19815), four animals had no viable library B bacteria; in these animals library B DNA tags were only found in homogenates of sterile granulomas (Supp. Table 3). Only 1 of the 9 granulomas with both library A and B DNA tags grew library B bacteria. Thus, primary infection fully protected 5 out of 8 animals against productive (CFU+) infection with the challenge strain (**Fig. 2A).** Collectively, these data suggest that primary Mtb infection initiates an immune response which leads to rapid neutralization of the challenge strain.

**Figure 2:**
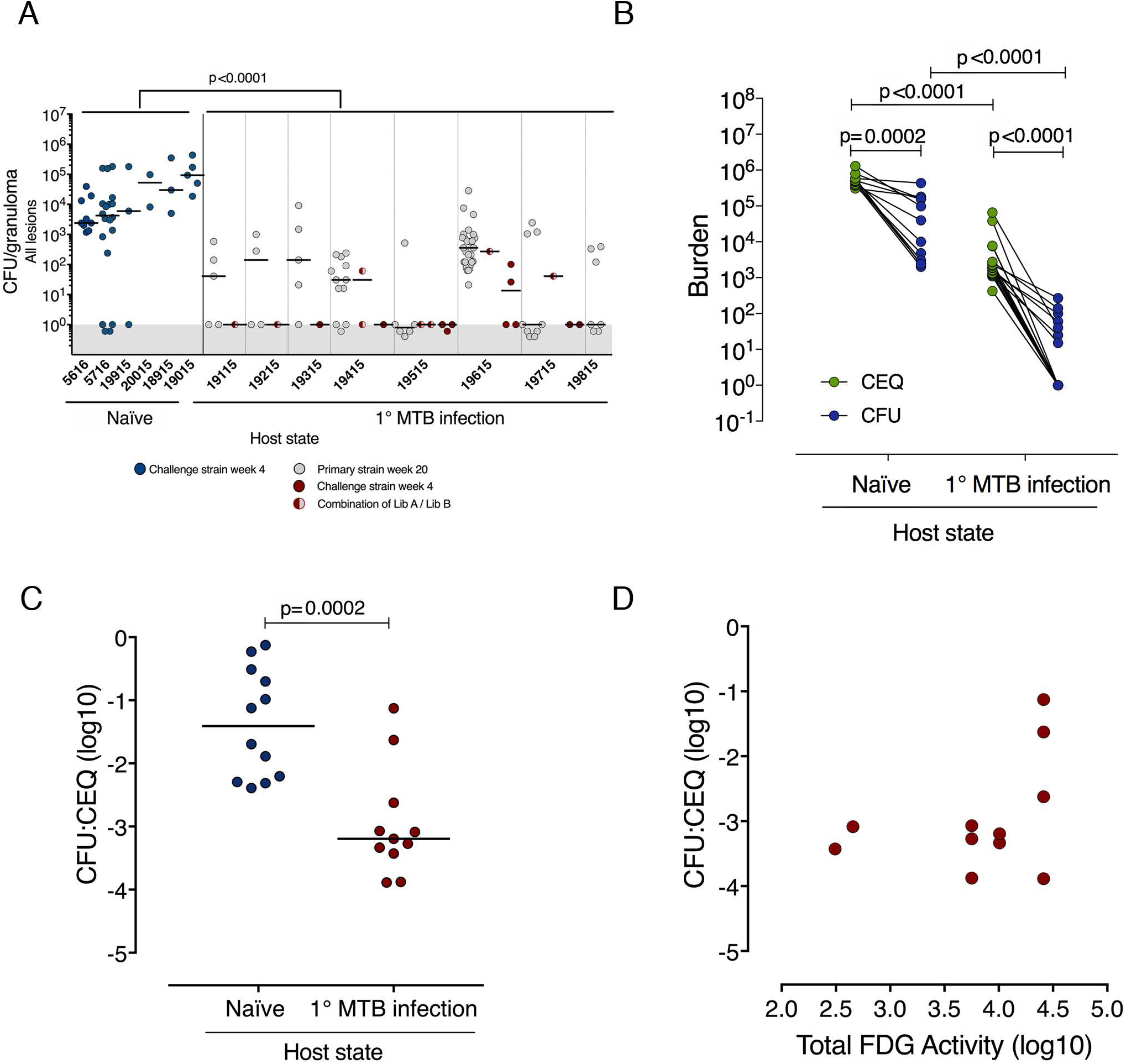
Reinfection promotes sterilizing immunity in secondary strain granulomas and restricts thoracic lymph node dissemination. (a) The bacterial burdens (CFU) of all granulomas in naïve macaques (blue), or reinfection macaques (Library A: gray; Library B: red; Library A and B: red/gray circles) 4 weeks post-Mtb Library B challenge. The CFU for all granulomas in the reinfected macaques were significantly reduced relative to the naïve monkeys. N=42 ranging from 2 to 20 per animal (naïve) and N=94 ranging from 5 to 32 per animal (reinfection). Each symbol is a granuloma, p < .0001, Mann-Whitney test. (b) Total bacterial genome counts (CEQ) and viable bacterial burden counts (CFU) were determined from the same Library B granulomas from naïve or reinfection macaques and bacterial killing (CFU/CEQ) for each granuloma shown in (c), p = 0.0002, Mann-Whitney test. Each symbol for (b) and (c) is a granuloma. CFU was significantly lower than CEQ in both naïve and reinfection granulomas (b), paired t-tests, p-values reported in figure. Both CEQ and CFU were reduced in reinfection granulomas compared with granulomas from naïve animals (b), Mann-Whitney tests, p-values reported in figure. (d) There was no correlation between extent of host lung inflammation (total lung FDG activity) and bacterial killing (CFU/CEQ) of library B granulomas (Spearman r = 0.38, p = 0.2454). In b, N=12 (naïve) and N=21 (reinfection). In b and c, N=12 (naïve) and N=11 (reinfection).

Note that the bacterial loads in granulomas containing library A, delivered 20 weeks prior to necropsy, were also lower than library B granulomas from naïve animals analyzed ∽4 weeks post infection. However, this decrease in bacterial load over time is consistent with our previously published studies^12^ and data from historical macaques infected with Mtb Erdman (Supp. Fig. 2A), where CFU/granuloma decreases after the onset of adaptive immunity, and did not support the hypothesis that rechallenge altered the course of the primary infection. To further investigate this question, we assessed inflammation in individual library A granulomas by PET CT prior to and after library B challenge and compared this to similar time points in historical macaques challenged with Mtb Erdman, and again saw no evidence that secondary infection altered the inflammatory dynamics of the preexisting infection (Supp. Fig. 3).

**Figure 3:**
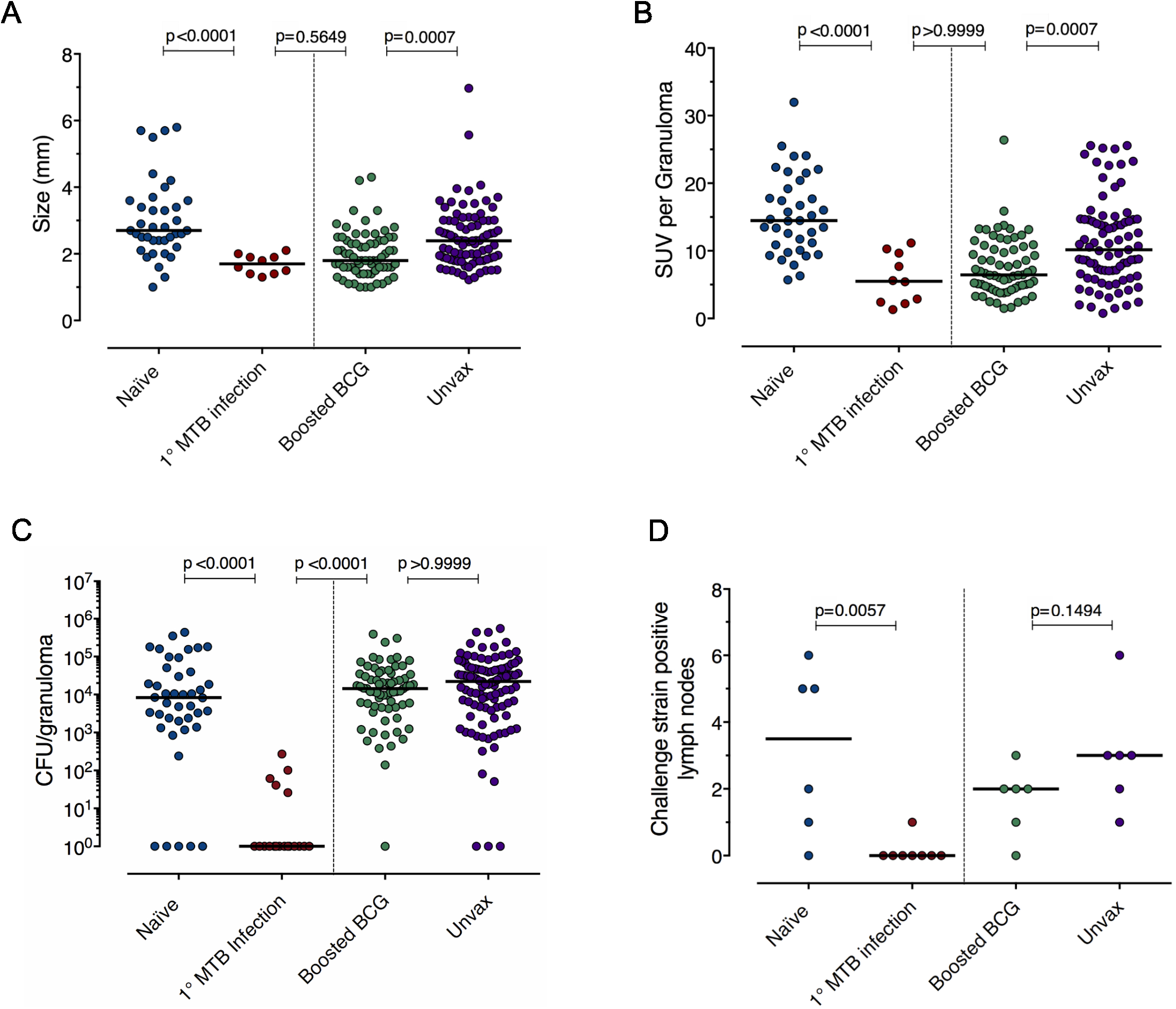
Primary Mtb infection reduces bacterial burden in secondary challenge granulomas, while boosted BCG has no effect on early granuloma bacterial burden. (a, b) Size (mm, by CT) and inflammation (as measured by [^18^F]-FDG avidity via standard uptake value, SUV) for Library B (for naïve and reinfection macaques) or Mtb (for BCG+H56 and unvaccinated controls) granulomas. (c) CFU of Mtb Library B granulomas from naïve (blue) and reinfection (red) macaques and 5-6 week granuloma CFU in boosted BCG (green) and unvaccinated control (purple) macaques challenged with Mtb Erdman. For a, b, and c, each symbol is a granuloma, p < 0.0001, Kruskal-Wallis tests, Dunn’s multiple comparison adjustment was used and adjusted p-values are reported in the figure above corresponding comparisons. All granulomas from N=6 macaques (naïve); N=7 macaques (reinfection); N=6 BCG+H56 macaques; N=6 unvaccinated macaques are represented. (d) Left panel: Number of thoracic lymph nodes from each macaque from which were CFU+ for Mtb Library B. Right panel: Number of thoracic lymph nodes from unvaccinated or BCG+H56 vaccinated CFU+ positive for Mtb at 5-6 weeks post-infection. N=6 (naïve), N=8 (reinfection), and N=6 (unvaccinated). Each symbol is a macaque, Mann-Whitney test p-values reported in figure.

We next sought to distinguish whether the lower bacterial burden in granulomas formed after reinfection reflected restriction of bacterial growth and/or true enhancement of bacterial killing. We assessed granuloma bacterial genome counts (CEQ) attributable to library B in naïve and rechallenged animals and then quantified killing of the challenge strain by relating viable library B live bacteria counts (CFU) to library B genome counts (CEQ), as described^12,22^ (**Fig. 2B**). In rechallenged animals, library B granulomas had significantly lower CEQ than in naïve animals, indicating reduced replication of the infecting bacteria. Library B granulomas had concomitantly lower CFU (**Fig. 2B**), which also reflected significantly increased killing (as reflected by CFU/CEQ) of the library B in granulomas from rechallenged animals relative to naïve animals (**Fig. 2B, C** median log killing = -1.41 (naïve) vs. -3.19 (reinfected), p=0.0002; ∽1.75 log increase in killing). Interestingly, there was no correlation between bacterial killing of library B Mtb and host lung inflammation at the time of second challenge (**Fig. 2D**, r = 0.03, p = 0.8832), nor was there any association between total thoracic bacterial burden at necropsy in those macaques who had few library B+ granulomas and those who had none (Supp. Fig 2B). Thus, the extent of disease at time of secondary exposure does not appear to affect the protection against reinfection.

To assess the protection provided by concurrent infection in the context of current vaccine strategies, we compared the bacterial loads from granulomas in our naïve and reinfection monkeys to those from macaques vaccinated with BCG boosted with an adjuvanted fusion protein (H56 in CAF01^23,24^). We previously showed that BCG+H56 provides protection against reactivation, reduces pathology and improves survival of Mtb-infected macaques^25,26^. Moreover, the size and FDG avidity of early granulomas in BCG+H56 vaccinated animals were reduced compared to unvaccinated animals, as was found in the granulomas formed in the setting of reinfection (**Fig. 3A, B**). However, at the early time points assessed here, the bacterial control engendered by primary Mtb infection was dramatically superior to that found in the vaccinated animals (**Fig. 3C**). In addition, primary infection prevented dissemination of the reinfection strain to thoracic lymph nodes while BCG+H56 did not have a similar effect (**Fig. 3D**); the number of granulomas at 4-5 weeks post-challenge in BCG-vaccinated macaques was similar to contemporaneous control macaques. In aggregate, primary infection provided ∽10,000 fold protection against Mtb reinfection as assessed by granuloma bacterial burdens (Fig. 3C), reflecting the combined effects of restricted establishment of infection, limited bacterial growth, and increased bacterial killing. These data suggest that there is a fundamental difference in the immune profile elicited by primary Mtb infection from that generated by BCG intradermal vaccination, most likely reflecting the difference in immunity expressed by local resident T effector cells maintained by ongoing infection versus the systemic response promoted by a distal boosted BCG vaccination.

To provide insight into the local immunologic landscape that may contribute to the protection seen with reinfection, we first assessed uninvolved lung tissue from reinfected monkeys using Luminex (Supp. Fig. 4A) compared to responses in an uninfected monkey, where it was only possible to euthanize one animal for analysis of normal lung tissue. These data suggested that even in uninvolved tissue, infection is associated with the expression of innate cytokines and chemokines. Though this analysis was limited by the availability of uninfected animals, the data suggest increased IL-8 in lung tissue from infected monkeys compared to uninfected control monkeys, possibly implicating increased neutrophil migration in infected tissues, and a decrease in IL-1 in infected tissues, with a trend towards increased IL-1RA. CXCL9, a chemokine that binds CXCR3 was also slightly higher in infected lung tissues, suggesting possible increased T cell migration, while CXCL13, which is chemotactic for B cells, was reduced. We also assessed uninvolved lung tissue from a separate set of Mtb-infected monkeys (not reinfected) at similar time points for Mtb-specific (ESAT6/CFP10) T cell responses (Supp. Fig. 4B). Mtb-specific T cell responses were higher in infected lung tissue, compared to uninfected lung tissue, indicating the presence of T cells resident within lung tissue that could rapidly activate macrophages upon encountering a new Mtb bacillus. Thus, a reinfecting bacillus encounters innate and adaptive immune responses in lung tissue, which may underlie the more rapid killing of the bacterium that we observed.

**Figure 4:**
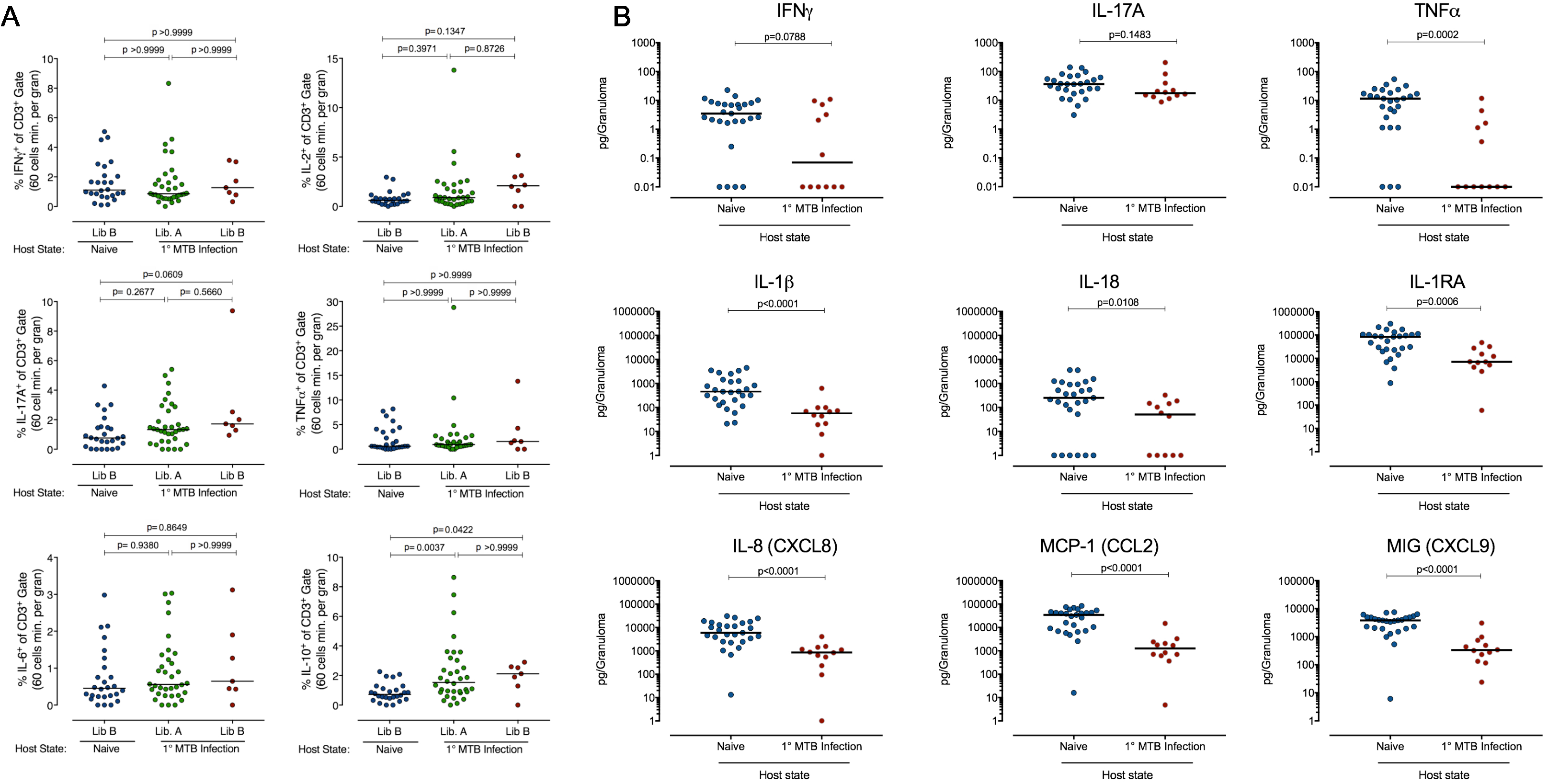
Reinfection challenge strain granulomas have elevated IL-10 T cell responses and reduced levels of pro-inflammatory cytokines and chemokines. (a) Granuloma T cell responses by intracellular cytokine staining and flow cytometry from naïve and reinfection animals. N=26 (naïve), N=34 (Library A, reinfection), and N=7 (Library B, reinfection). Each dot is a granuloma, Kruskal-Wallis test, Dunn’s multiple comparison adjustment was used and adjusted p-values are reported. (b) Supernatants from library B granulomas from naïve or reinfection macaques were assessed for cytokine and chemokines. N=27 (naïve) and N=12 (reinfection). Each dot is a granuloma, p < 0.05-0.0001, Mann-Whitney test.

We next assessed T cell responses in the few library B granulomas that did establish in the reinfected monkeys and compared these to library A granuloma in the same animals and library B granulomas that established in naïve animals (Fig. 4A). In general, T cell responses in library A and B granulomas in reinfected macaques were similar, and the data reflect the normal variability seen in immune responses in granulomas^18^. There were no significant differences in the frequency of T cells making T_H_1 cytokines among library B granulomas from naïve or reinfected macaques, or library A granulomas in reinfected macaques. However, IL-10 was significantly higher in both library A and B granulomas in reinfected macaques compared to granulomas in naïve macaques. Our prior data suggested that the combination of T cells expressing IL-10 and T cells expressing a pro-inflammatory cytokine in granulomas was associated with the sterilization of Mtb granulomas^18^. The higher levels of IL-10 in granulomas with lower bacterial loads as compared to granulomas in naïve animals is consistent with this finding.

More broadly, production of inflammatory cytokine and chemokines (measured in granuloma supernatants) was lower in library B granulomas in animals with primary infection relative to those formed in naïve animals (**Fig. 4D**). TNF, IL-1β, IL-18, IL-1RA, IL-8, MCP-1, and MIG were all present at significantly (p<0.05-0.0001) lower levels in secondary granulomas, with a trend towards lower IFN-γ. This likely reflects the low bacterial burden in the reinfection granulomas and again may indicate a healing response as those granulomas rapidly kill the bacteria. Further study into the precise kinetics and cellular organization of these granulomas is needed to help inform intervention strategies.

Overall, we present evidence for robust concomitant immunity in the cynomolgus macaque model of TB. We found complete protection against productive secondary infection in five macaques with ongoing infection (**Fig. 1D, 2A**), almost no productive dissemination to lymph nodes (**Fig. 3D**), and a ∽10,000-fold decrease in live Mtb in library B (reinfection) granulomas compared to age-matched granulomas in naïve animals (**Fig. 2B,C, Fig. 3C**). One monkey had no trace of library B DNA despite receiving a secondary challenge dose of 10 CFU. This level of protection would be truly remarkable for a TB vaccine, and suggests that a protective vaccine for TB may have to induce immune responses even greater than those induced by primary infection. The recent study showing substantial protection in rhesus macaques by a CMV vector expressing multiple Mtb antigens suggests that sustained high level T cell responses are important for immunity but the correlate of protection analysis also pointed to the potential importance of an activated innate immune response in mediating bacterial clearance^27^. We could not formally define a correlate of protection conferred by concurrent infection since all concurrently infected animals were robustly protected against reinfection; there were no immunologic or disease parameters that segregated animals that fully sterilized the challenge from those that sterilized many but not all sites of infection (Supp. Fig. 5). However, our immunologic analysis does provide evidence of an activated innate response and *Mtb* specific T cells in the uninvolved lung tissue of infected animals that could participate in clearance.

Concomitant immunity has been described in other systems, e.g. tumor rejection models^28^. In terms of infectious diseases, this is best characterized for leishmaniasis, where the presence of a live but contained *Leishmania* infection prevents establishment of a new infection. In the Leishmania model, this protection depends on the presence of high levels of T effector cells rather than central memory T cells, and is lost when the original infection is cleared^29^. Our study supports concomitant immunity in Mtb infections, although it remains to be determined whether complete clearance of primary Mtb infection with drug treatment will abrogate the extraordinary protective effects shown here^30^. While the precise immune mechanisms underlying the protection described here are not yet fully characterized, this study provides exciting clues for revised TB vaccine strategies to achieve both sterilizing immunity and protection from infection.

## Methods

### Macaque infections, PET-CT imaging, and tissue excision

Fourteen adult cynomolgus macaques (*Macacca fasicularis*) were obtained from Valley Biosystems (Sacramento, California) and screened for Mtb and other comorbidities during a month-long quarantine. Each macaque had a baseline blood count and chemical profile and was housed according to the standards listed in the Animal Welfare Act and the Guide for the Care and Use of Laboratory Animals. All procedures were approved by the Institutional Animal Care and Use Committee at the University of Pittsburgh. The animals were separated into two cohorts: 8 macaques were assigned to reinfection and 6 were assigned to naïve 4-week only controls. All animals were infected with barcoded strain Erdman Mtb via bronchoscopic instillation as previously published^9,10^. The infection schema is provided in **Fig. 1A**; 8 macaques were infected with Mtb library A, followed 16 weeks later by Mtb library B; 6 naïve macaques were infected with Mtb library B in a series of matched infections. All animals received an inoculum of <15 CFU (determined by plating a sample of the inoculum and counting CFU after 3 weeks) with the details listed in **Supp. Table 1**. The animals were further subdivided such that 4 reinfection animals were directly paired with 2, naïve animals; 2 additional naïve animals were infected separately (**Supp. Table 1**). Each macaque was followed with serial 2-deoxy-2-[^18^F]-fluoro-D-glucose ([^18^F]-FDG) PET-CT imaging as previously described^12-14^ to identify and track lesion formation and progression over time. PET-CT scans were performed monthly during the primary infection, and as noted in Supplementary Table 1 after reinfection. Total FDG activity in lungs was used to estimate thoracic bacterial burden prior to reinfection, as previously published^11,31^. Granulomas were individually characterized by their date of establishment (scan date), size (mm), and relative metabolic activity as a proxy for inflammation ([^18^F]-FDG standard uptake normalized to muscle [SUVR]^11,31^). Granulomas ≥ 1mm can be discerned by our PET-CT imaging analysis (**Fig. 1B**).

All macaques were necropsied at ∽4 weeks post-library B infection. The pre-necropsy PET-CT scan was used to map new and old granulomas. To avoid barcode cross-contamination, individual granulomas were separately excised and processed, along with all thoracic lymph nodes. Each sample was homogenized to a single cell suspension, plated for CFU on 7H11 agar supplemented with oleic albumin dextrose catalase (OADC), and frozen aliquots were stored for DNA extraction.

For the BCG+H56 vaccine study, 6 cynomolgus macaques were vaccinated with BCG (5×10^5^ ID) followed by an intramuscular injection of the fusion protein H56 (composed of ESAT-6, Ag85B, and Rv2660c) in the adjuvant CAF01 at 10 and 14 weeks post-BCG. Six cynomolgus macaques were unvaccinated controls. Six months after BCG, all 12 macaques were infected with *M. tuberculosis* strain Erdman (not barcoded) (**Supp. Table 3**). PET-CT scans were performed at 4-5 weeks post-infection, just prior to necropsy (**Supp. Table 2**), and analyzed as in the reinfection study. All granulomas and lymph nodes were obtained using the PET-CT scan as a map, homogenized and plated for bacterial burden.

### Isolation and preparation of bacteria genomic DNA from tissue samples

A small portion of granuloma homogenate was frozen for qTag sequencing and chromosomal equivalent (CEQ) analysis. For analysis of bacteria that grew from the granulomas, bacterial colonies from granuloma and lymph node plates were scraped to obtain all colonies, and frozen. Genomic DNA was extracted as previously published^12,22^. In brief, gDNA was twice extracted with phenol:chloroform:isoamyl alcohol (25:24:1, Invitrogen) with an intermediate bead beating step using 0.1mm zirconia-silica beads (BioSpec Products, Inc.).

### Barcode/qtag determination via sequencing and nanostring

To identify DNA tags from each library, genomic DNA was diluted to 10 ng/μL and amplified with Q5 polymerase (New England Biolabs) with two rounds of PCR of 8-15 cycles each, using the primers described in Martin *et al*^15^. Samples were then sequenced on an Illumina MiSeq using v2 chemistry. Barcodes were identified using custom scripts as described in Martin *et al.* To identify qTags, genomic DNA was diluted to 100 ng/μL and amplified with 24-36 cycles of PCR, using the primers listed in Supplementary Table 3. PCR product was used as input in the NanoString nCounter assay (NanoString Technologies) with custom-designed probes to determine qTag identity. We attributed the bacteria in any mixed lesion to library B, a conservative assumption that would lead us to overestimate the success of library B in the setting of rechallenge.

### Luminex on granuloma supernatant and lung tissue

Supernatants from granuloma and uninvolved lung tissue homogenates were frozen at -80°C until time of assay. After thawing, samples were filtered using a 0.22um syringe filter to remove any infectious bacteria and kept on ice. Thirty cytokines and chemokines were assayed from the granuloma supernatants in duplicate using a ProcartaPlex multiplex immunoassay (Invitrogen) specific for nonhuman primate samples according to manufacturer’s instructions, with an additional dilution of the supplied standard curve to extend sample detection range. Multiplex results were read and analyzed by BioPlex reader (BioRad).

### T cell responses in granulomas and uninvolved lung tissue

Flow cytometry was performed on individual granulomas from naïve and reinfected macaques and a random sampling of lung tissue without any disease pathology from 6 Mtb infected macaques (with no reinfection) and an uninfected macaque. For granulomas, no restimulation was performed because of the high level of antigens in the granulomas, as previously noted^18^. Single cell suspension of the lung tissue was stimulated with peptide pools of Mtb specific antigens ESAT-6 and CFP-10 (10ug/ml of every peptide) in the presence of Brefeldin A (Golgiplug: BD biosciences) for 3.5 hours at 37°C with 5% CO_2_. The cells were then stained for Viability marker (Invitrogen), surface and intracellular cytokine markers. Flow cytometry for cell surface markers for T cells included CD3 (clone SP34-2; BD Pharmingen), CD4 (Clone L200, BD Horizon) and CD8 (clone SK1, BD biosciences). Intracellular cytokine staining panel included pro-inflammatory cytokines: Th1 [IFN-γ (Clone B27), IL-2 (Clone: MQ1-17H12), TNF (Clone: MAB11)] and Th17 [IL-17 (Clone: eBio64CAP17). Data acquisition was performed using an LSR II (BD) and analyzed using FlowJo Software v.9.7 (Treestar Inc, Ashland, OR).

### PBMC PCA analysis

A dimension reduction technique was performed on data extracted from PBMCs sampled just prior to reinfection (15 weeks post infection). Principal components analysis (PCA) was utilized here to investigate whether protected animals cluster separately from unprotected animals using blood signatures. Unfortunately, with such a small sample size (n = 8 macaques) PCA is not stable, so these results should be interpreted with caution.

### Statistical analysis

Graphs were created in Graphpad Prism and JMP and all statistical analysis was performed in Graphpad Prism. In datasets that included zeroes, data were transformed by adding either 1 or 0.01 so that zeroes could be graphed on a log-scale. The D’Agostino and Pearson test was used to test for normality. If data were found to be normal, groups were compared using t-tests; otherwise, groups were compared using the non-parametric Mann-Whitney test for two groups or the Kruskal-Wallis test for three groups (with Dunn’s multiple comparison adjustment). All tests were two-sided with significance defined as p<0.05.

## Acknowledgements

We thank the members, laboratory technicians and veterinary staff of the Flynn and Fortune laboratories for their technical expertise and assistance with animal care, sample processing, and study design. We are grateful to Dr. Barry Bloom for helpful discussions. This work was made possible by support from Aeras (JLF, SMF) and the National Institutes of Health, R01 AI114674 (JLF, SMF) and T32 AI089443 (AMC). Support was also provided by the Burroughs Wellcome Foundation (SMF), and the Harvard Center for AIDS Research P30 A1060354 (CJM, SMF).

## Supplementary Figure Legends

**Suppl. Figure 1:**
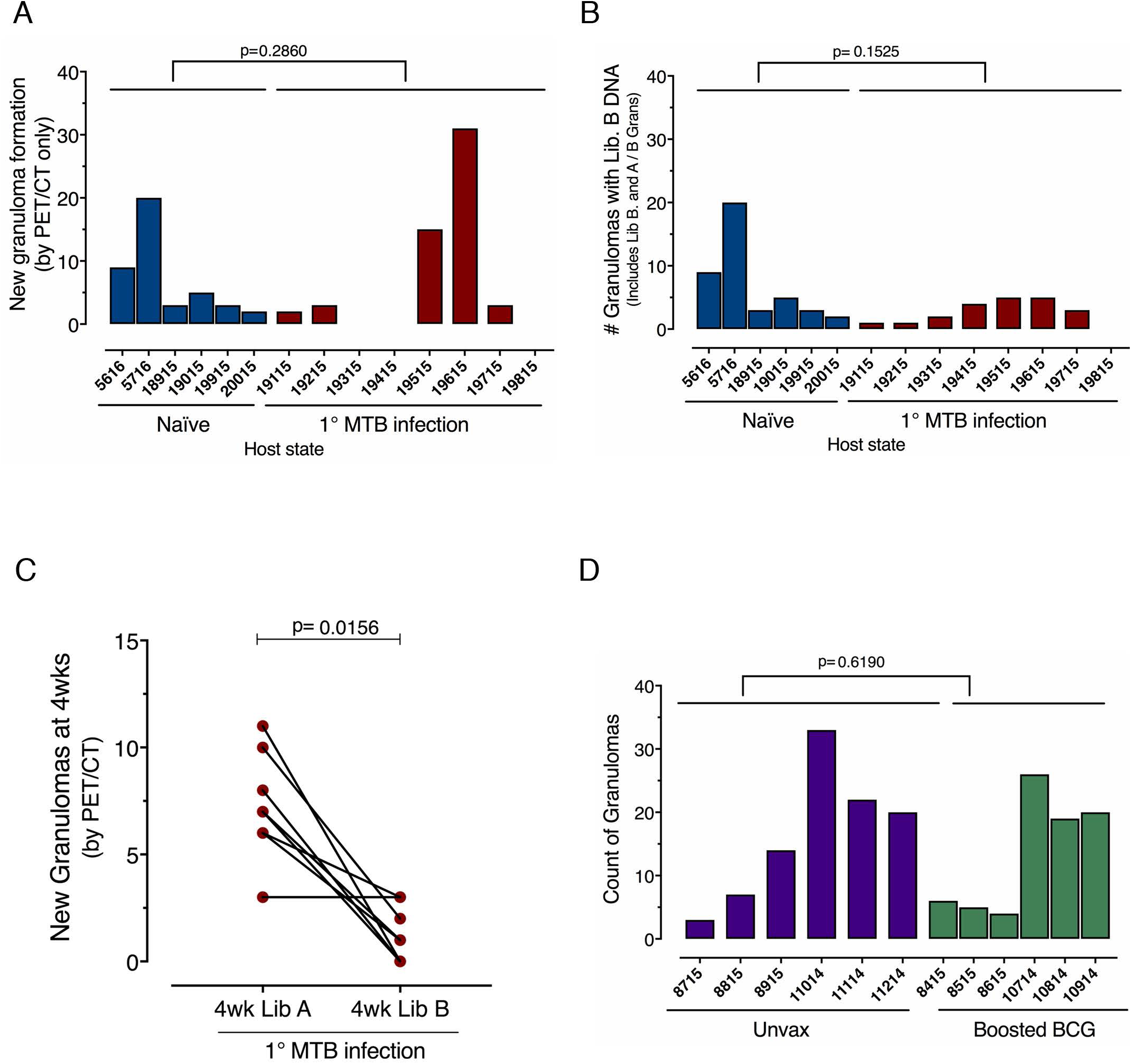
There is no difference in the total numbers of granulomas that contain library B in reinfection. (a) All new granulomas seen by PET CT in naïve and reinfected macaques at 4 weeks post-Library B infection. (b) All granulomas with Library B DNA in naïve or reinfected macaques. This count includes both newly established and pre-existing granulomas that contained library B in the reinfection cohort. Monkey ID: 19815 had no detectable library B in any tissue. (c) Number of new granulomas seen by PET CT established by library A or library B 4 weeks post-infection in the same animal. p = 0.0156, Wilcoxon-matched pairs signed rank test. (d) Number of granulomas seen by PET CT at 4-5 weeks post-Mtb Erdman infection between unvaccinated and BCG+H56 vaccinated macaques. Statistics: Mann-Whitney test.

**Suppl. Figure 2:**
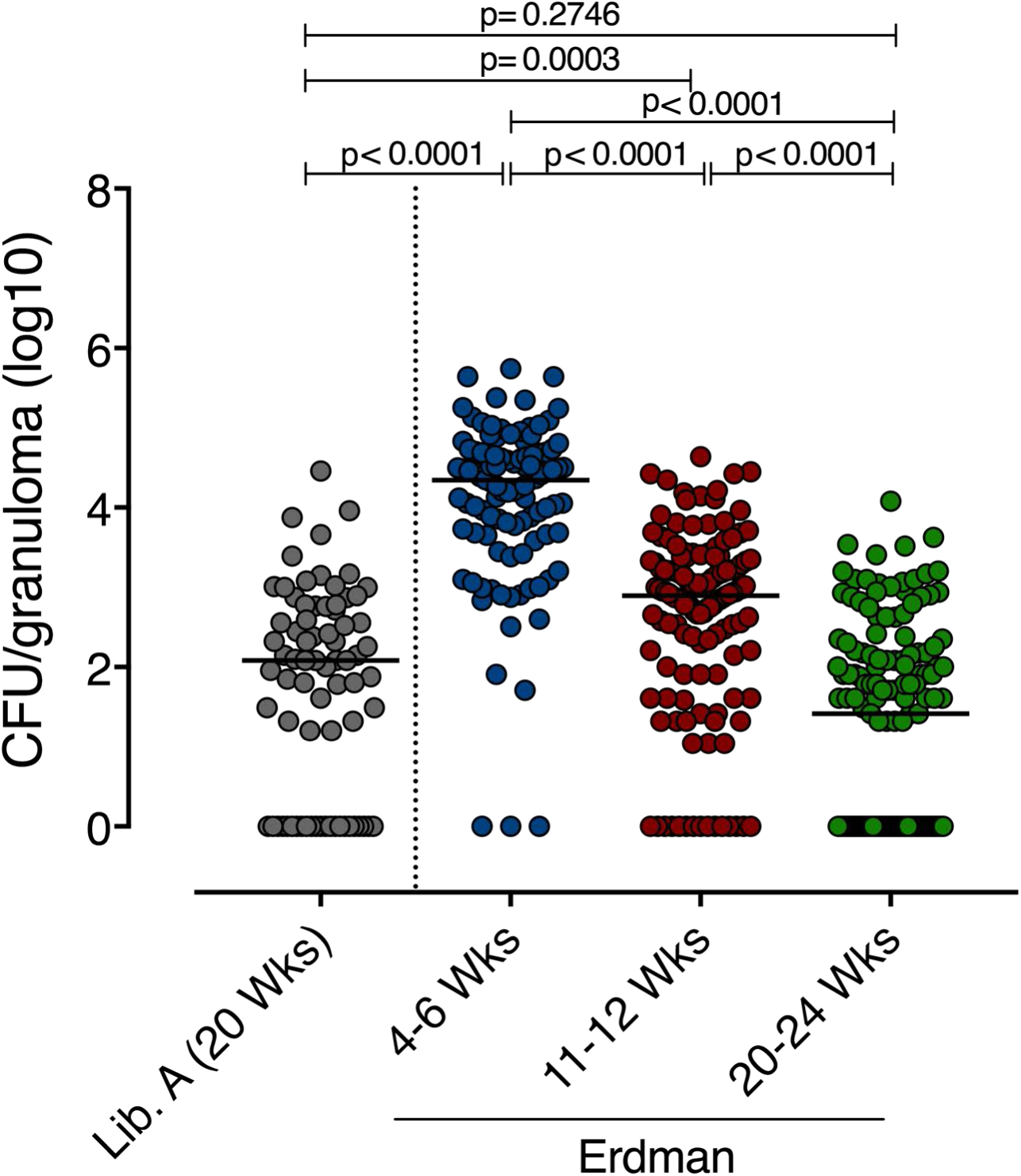
Bacterial burden of granulomas is reduced over time. CFU/granuloma was compared from Mtb Erdman infected macaques at 4-6, 11-12, and 20-24 weeks post-infection, using historical controls (N=17). This is similar to data published in Lin, Ford, et al, but using a different set of monkeys. On the left side of the graph, CFU/granuloma of library A in our reinfected animals is shown (N=8); these values are similar to the values of Erdman at 20-24 weeks.

**Suppl. Figure 3:**
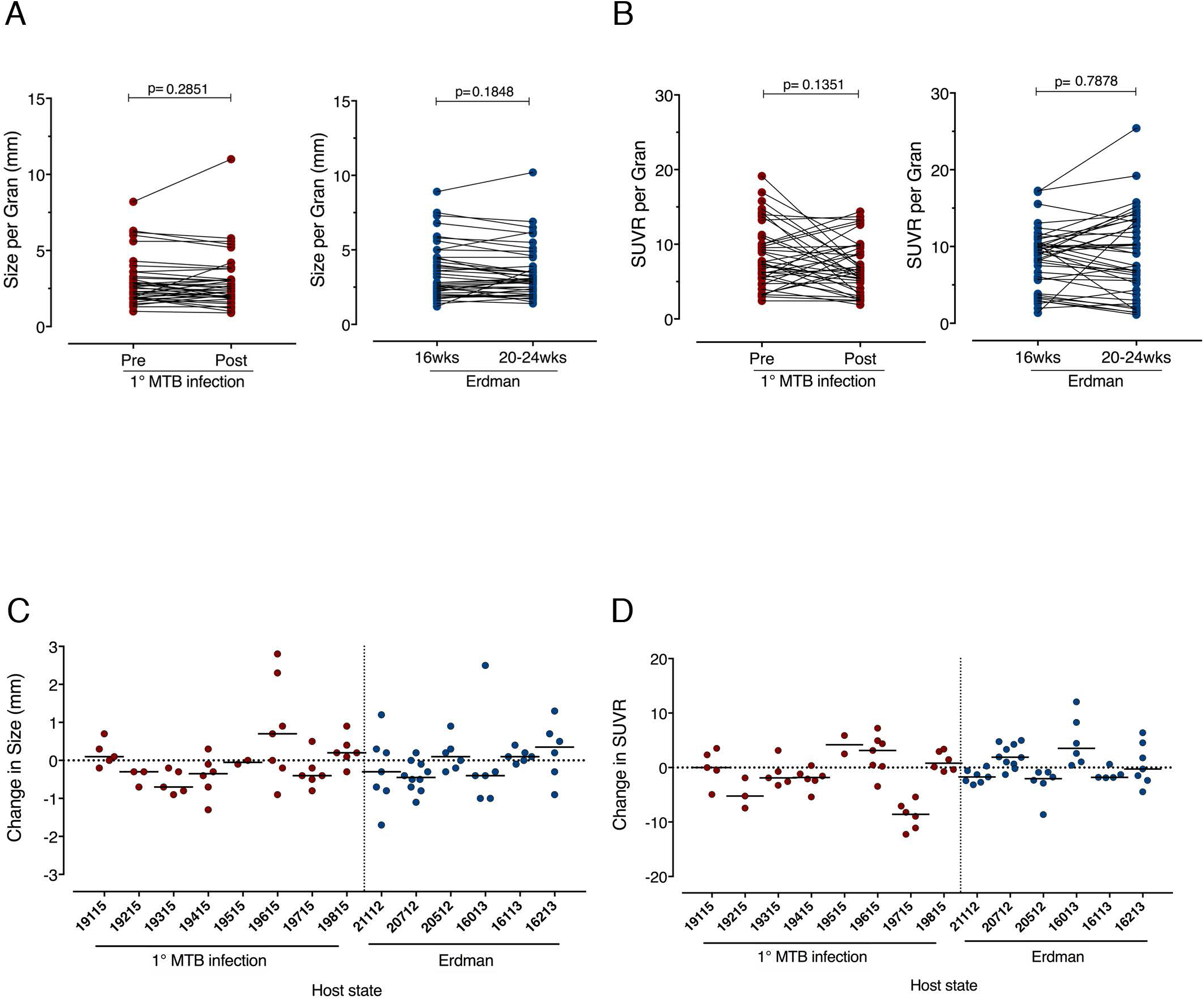
Size (in mm) and FDG activity (SUVR) of individual library A granulomas in animals pre- and post-Library B reinfection and historical controls from 16 – 20 weeks post-infection. (A) Granulomas do not change significantly in size after infection with Library B in the reinfection animals (p = 0.2851). Similarly, granulomas from animals infected with the Erdman strain do not change significantly in size from 16 to 20 (or 24) weeks post-infection (p = 0.1848). (B) Previously established (library A) granulomas do not significantly increase or decrease in FDG activity (SUVR) after reinfection with Library B (p = 0.1351). Likewise, in Erdman-infected historical controls, granulomas do not significantly change in SUVR (p = 0.7878) from 16 to 20 weeks post-infection. (C) Change in size (mm) per granuloma by animal. (D) Change in FDG activity (SUVR) per granuloma by animal. Each symbol represents a granuloma. Statistics: Wilcoxon matched-pairs signed rank test.

**Suppl. Figure 4:**
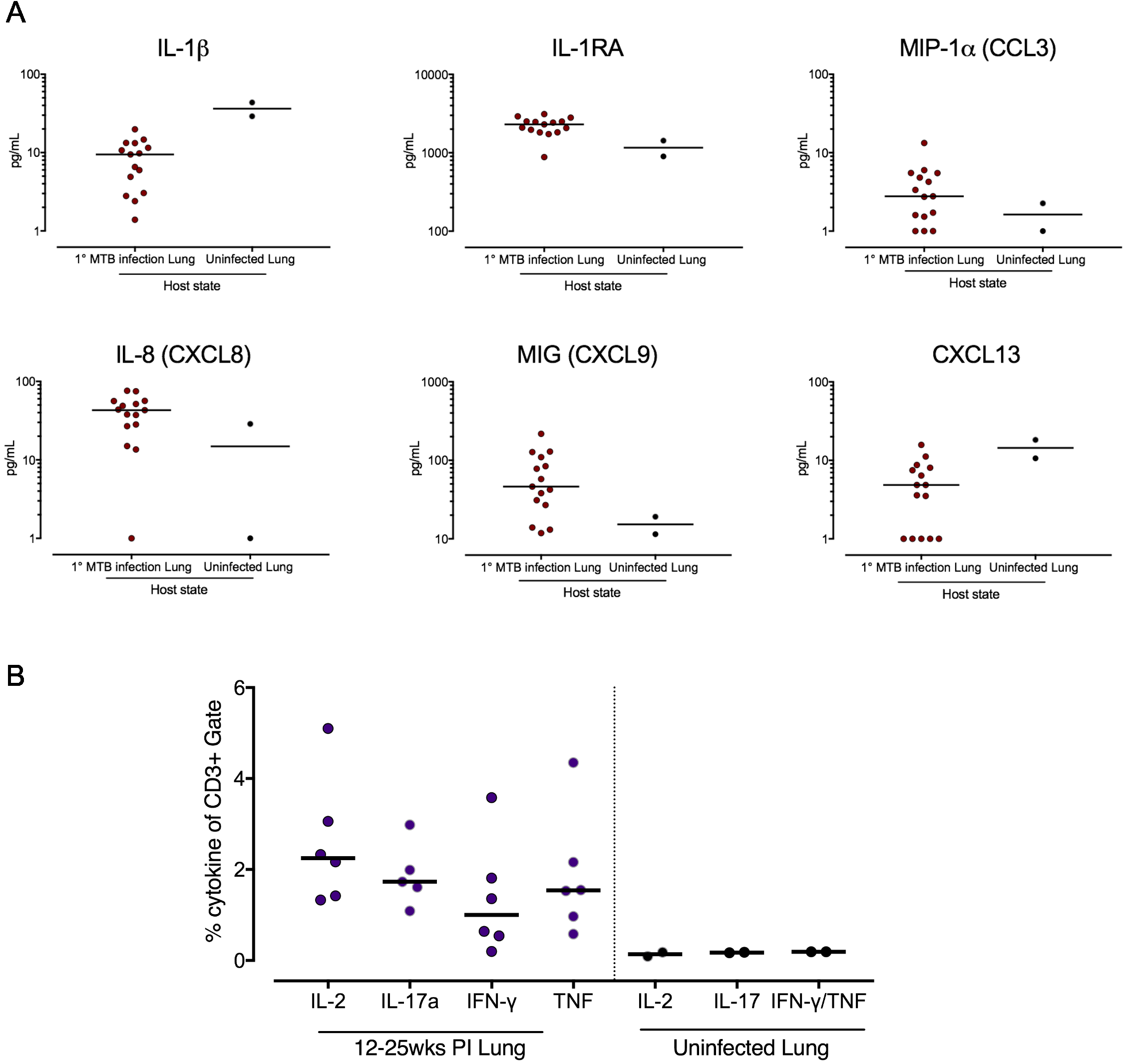
Uninvolved lung tissue from infected macaques has increased levels of chemokines and higher Mtb-specific T cell responses compared to lung tissue from an uninfected macaque. A. Luminex analysis on supernatant from uninvolved (no granuloma) lung tissue was compared between reinfected macaques and an uninfected macaque. B. Using a separate set of macaque lung tissue (20-24 weeks post-infection but not reinfected), the T cell responses following ESAT-6 and CFP10 stimulation were assessed by flow cytometry, and compared to an uninfected macaque. No statistics were performed, due to the small sample size for the uninfected macaque lung.

**Suppl. Figure 5:**
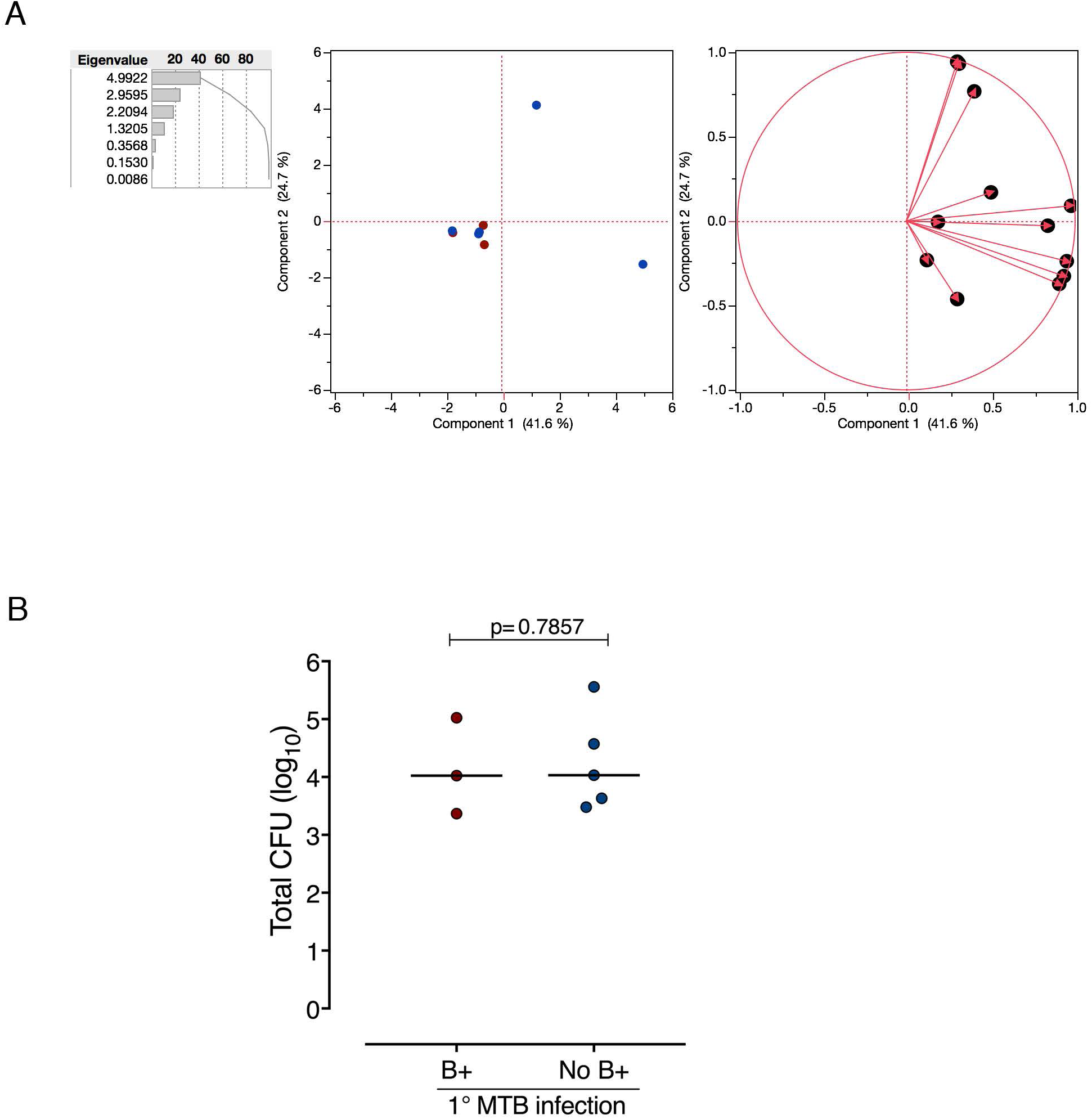
(A) Principal Components Analysis of PBMC data. Far left is the Eigenvalue Pareto Plot showing the cumulative percentage of variation accounted for in each principal component. Middle is a scatterplot of the first two components color-coded for animals with CFU-positive B lesions (blue), and those without (red). There is no obvious clustering of these groups. Far right is a loading plot showing the correlations of the original variables to the first two principal components. B. Total thoracic CFU (log_10_) is similar between animals with and without CFU-positive Library B granulomas (p = 0.7857, Mann-Whitney test).

## References

1. Andrews, J.R., et al. Risk of progression to active tuberculosis following reinfection with Mycobacterium tuberculosis. Clinical infectious diseases: an official publication of the Infectious Diseases Society of America 54, 784–791 (2012).

2. Wang, J.Y., et al. Prediction of the tuberculosis reinfection proportion from the local incidence. J Infect Dis 196, 281–288 (2007).

3. Cohen, T. & Murray, M. Incident tuberculosis among recent US immigrants and exogenous reinfection. Emerg Infect Dis 11, 725–728 (2005).

4. Cohen, T., et al. Mixed-strain mycobacterium tuberculosis infections and the implications for tuberculosis treatment and control. Clin Microbiol Rev 25, 708–719 (2012).

5. Charalambous, S., et al. Contribution of reinfection to recurrent tuberculosis in South African gold miners. Int J Tuberc Lung Dis 12, 942–948 (2008).

6. Caminero, J.A., et al. Exogenous reinfection with tuberculosis on a European island with a moderate incidence of disease. American journal of respiratory and critical care medicine 163, 717–720 (2001).

7. Henao-Tamayo, M., et al. A mouse model of tuberculosis reinfection. Tuberculosis (Edinb) 92, 211–217 (2012).

8. Repique, C.J., Li, A., Collins, F.M. & Morris, S.L. DNA immunization in a mouse model of latent tuberculosis: effect of DNA vaccination on reactivation of disease and on reinfection with a secondary challenge. Infection and immunity 70, 3318–3323 (2002).

9. Capuano, S.V., et al. Experimental Mycobacterium tuberculosis Infection of Cynomolgus Macaques Closely Resembles the Various Manifestations of Human M. tuberculosis Infection. Infection and immunity 71, 5831–5844 (2003).

10. Lin, P.L., et al. Quantitative comparison of active and latent tuberculosis in the cynomolgus macaque model. Infection and immunity 77, 4631–4642 (2009).

11. Maiello, P., et al. Rhesus macaques are more susceptible to progressive tuberculosis than cynomolgus macaques: A quantitative comparison. Infection and immunity (2017).

12. Lin, P.L., et al. Sterilization of granulomas is common in active and latent tuberculosis despite within-host variability in bacterial killing. Nature medicine 20, 75–79 (2014).

13. Coleman, M.T., et al. Early Changes by (18)Fluorodeoxyglucose positron emission tomography coregistered with computed tomography predict outcome after Mycobacterium tuberculosis infection in cynomolgus macaques. Infection and immunity 82, 2400–2404 (2014).

14. Coleman, M.T., et al. PET/CT imaging reveals a therapeutic response to oxazolidinones in macaques and humans with tuberculosis. Sci Transl Med 6, 265ra167 (2014).

15. Martin, C.J., et al. Digitally Barcoding Mycobacterium tuberculosis Reveals In Vivo Infection Dynamics in the Macaque Model of Tuberculosis. mBio 8(2017).

16. Blumenthal, A., Trujillo, C., Ehrt, S. & Schnappinger, D. Simultaneous analysis of multiple Mycobacterium tuberculosis knockdown mutants in vitro and in vivo. PloS one 5, e15667 (2010).

17. Cadena, A.M., Fortune, S.M. & Flynn, J.L. Heterogeneity in tuberculosis. Nature reviews. Immunology 17, 691–702 (2017).

18. Gideon, H.P., et al. Variability in tuberculosis granuloma T cell responses exists, but a balance of pro- and anti-inflammatory cytokines is associated with sterilization. PLoS pathogens 11, e1004603 (2015).

19. Cadena, A.M., Flynn, J.L. & Fortune, S.M. The Importance of First Impressions: Early Events in Mycobacterium tuberculosis Infection Influence Outcome. mBio 7, e00342–00316 (2016).

20. Cosma, C.L., Humbert, O. & Ramakrishnan, L. Superinfecting mycobacteria home to established tuberculous granulomas. Nature immunology 5, 828–835 (2004).

21. Cosma, C.L., Humbert, O., Sherman, D.R. & Ramakrishnan, L. Trafficking of superinfecting Mycobacterium organisms into established granulomas occurs in mammals and is independent of the Erp and ESX-1 mycobacterial virulence loci. The Journal of infectious diseases 198, 1851–1855 (2008).

22. Lin, P.L., et al. PET CT Identifies Reactivation Risk in Cynomolgus Macaques with Latent M. tuberculosis. PLoS pathogens 12, e1005739 (2016).

23. Luabeya, A.K., et al. First-in-human trial of the post-exposure tuberculosis vaccine H56:IC31 in Mycobacterium tuberculosis infected and non-infected healthy adults. Vaccine 33, 4130–4140 (2015).

24. Aagaard, C., et al. A multistage tuberculosis vaccine that confers efficient protection before and after exposure. Nature medicine 17, 189–194 (2011).

25. Lin, P.L., et al. The multistage vaccine H56 boosts the effects of BCG to protect cynomolgus macaques against active tuberculosis and reactivation of latent Mycobacterium tuberculosis infection. The Journal of clinical investigation 122, 303–314 (2012).

26. Billeskov, R., et al. Testing the H56 Vaccine Delivered in 4 Different Adjuvants as a BCG-Booster in a Non-Human Primate Model of Tuberculosis. PloS one 11, e0161217 (2016).

27. Hansen, S.G., et al. Prevention of tuberculosis in rhesus macaques by a cytomegalovirus-based vaccine. Nature medicine (2018).

28. Gershon, R.K., Carter, R.L. & Kondo, K. On concomitant immunity in tumour-bearing hamsters. Nature 213, 674–676 (1967).

29. Peters, N.C., et al. Chronic parasitic infection maintains high frequencies of short-lived Ly6C+CD4+ effector T cells that are required for protection against re-infection. PLoS pathogens 10, e1004538 (2014).

30. Holmes, K.K., Bertozzi, S., Bloom, B.R. & Jha, P. (eds.). Disease Control Priorities, Third Edition: Volume 6. Major Infectious Diseases, (World Bank, Washington, DC, 2017).

31. White, A.G., et al. Analysis of 18FDG PET/CT Imaging as a Tool for Studying Mycobacterium tuberculosis Infection and Treatment in Non-human Primates. J Vis Exp (2017).

